# An RNAi screen of Rab GTPase genes in *C. elegans* reveals that somatic cells of the reproductive system depend on *rab-1* for morphogenesis but not stem cell niche maintenance

**DOI:** 10.1101/2024.12.03.626641

**Authors:** Kayt Scott, Noor Singh, Kacy Lynn Gordon

## Abstract

Membrane trafficking is a crucial function of all cells and is regulated at multiple levels from vesicle formation, packaging, and localization to fusion, exocytosis, and endocytosis. Rab GTPase proteins are core regulators of eukaryotic membrane trafficking, but developmental roles of specific Rab GTPases are less well characterized, potentially because of their essentiality for basic cellular function. *C. elegans* gonad development entails the coordination of cell growth, proliferation, and migration—processes in which membrane trafficking is known to be required. Here we take an organ-focused approach to Rab GTPase function *in vivo* to assess the roles of Rab genes in reproductive system development. We performed a whole-body RNAi screen of the entire Rab family in *C. elegans* to uncover Rabs essential for gonad development. Notable gonad defects resulted from RNAi knockdown of *rab-1,* the key regulator of ER-Golgi trafficking. We then examined the effects of tissue-specific RNAi knockdown of *rab-1* in somatic reproductive system and germline cells. We interrogated the dual functions of the distal tip cell (DTC) as both a leader cell of gonad organogenesis and the germline stem cell niche. We find that *rab-1* functions cell-autonomously and non-cell-autonomously to regulate both somatic gonad and germline development. Gonad migration, elongation, and gamete differentiation—but surprisingly not germline stem niche function—are highly sensitive to *rab-1* RNAi.

**SUMMARY:** The Rab family of GTPases regulate vesicular trafficking in cells. This study assessed the consequences for the growth of the gonad of RNAi-mediated gene knockdown of all Rab GTPase genes in *C. elegans*. The highly conserved primary regulator of ER-Golgi trafficking, *rab-1*, is essential for normal gonad and germline development. Further experiments found that *rab-1* is required in the somatic gonad for gonad elongation and migration, germline proliferation, and proper gamete formation. Surprisingly, the ability of the germline stem cell niche to maintain germ cells in the proliferative stem-like state was not affected by *rab-1* RNAi knockdown.

## INTRODUCTION

Members of the Rab family of GTPases—within the Ras superfamily—broadly regulate vesicular traffic within eukaryotic cells, including in specialized cell functions like cell division (Gibieža and Prekeris, 2018), cell polarity (Parker et al., 2018), and the release of neuropeptides into the synapse (Sasidharan et al., 2012). When systematically knocked out in a cell culture system with a method that is sensitive to potential redundancy among paralogs, specific functions for Rab family members have been revealed (Homma et al., 2019). While Rab GTPase-dependent membrane trafficking has been studied in *C. elegans* (Sato et al. 2014), only a handful of the 31 members of the Rab family have been investigated at a developmental genetic level (Ghosh and Sternberg, 2014; Lundquist, 2006). We hypothesize that the absence of literature exploring the roles of many Rab genes in *C. elegans* development is due to both redundancy among members of this family and essentiality of many Rab family members; nearly half of the Rab genes have been shown to cause lethality with strong loss of function in *C. elegans* (Table 1).

**Table 1.**
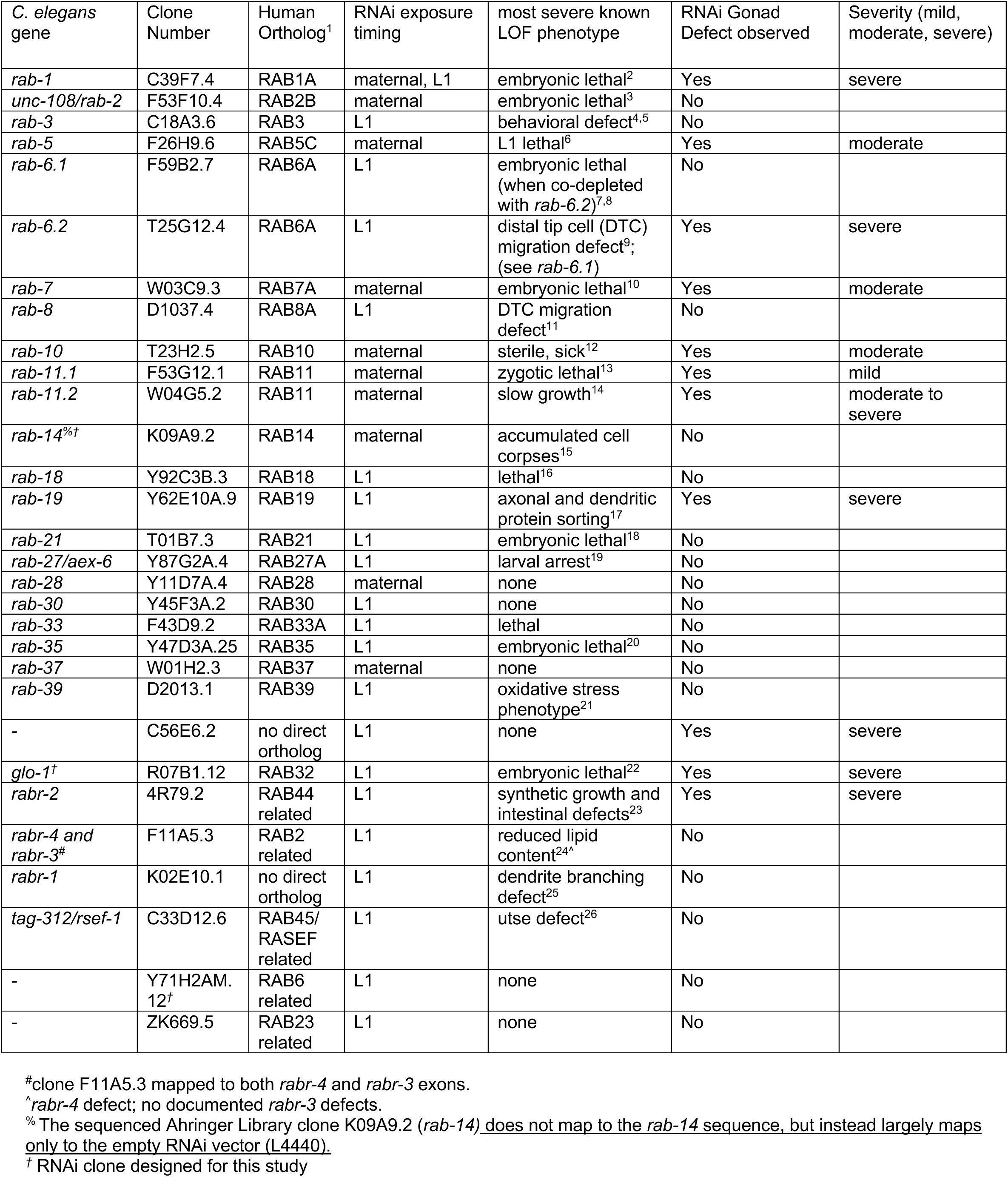
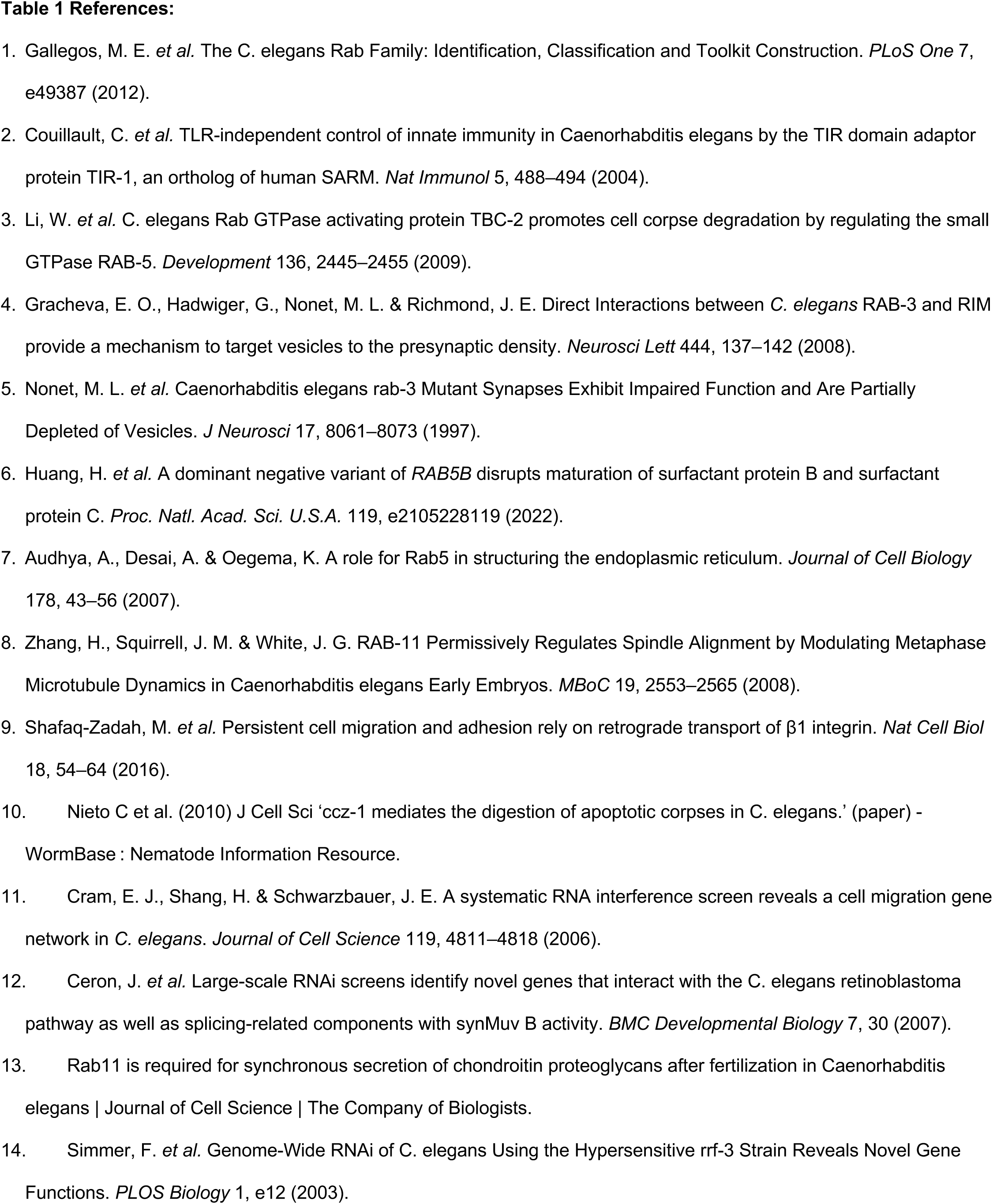
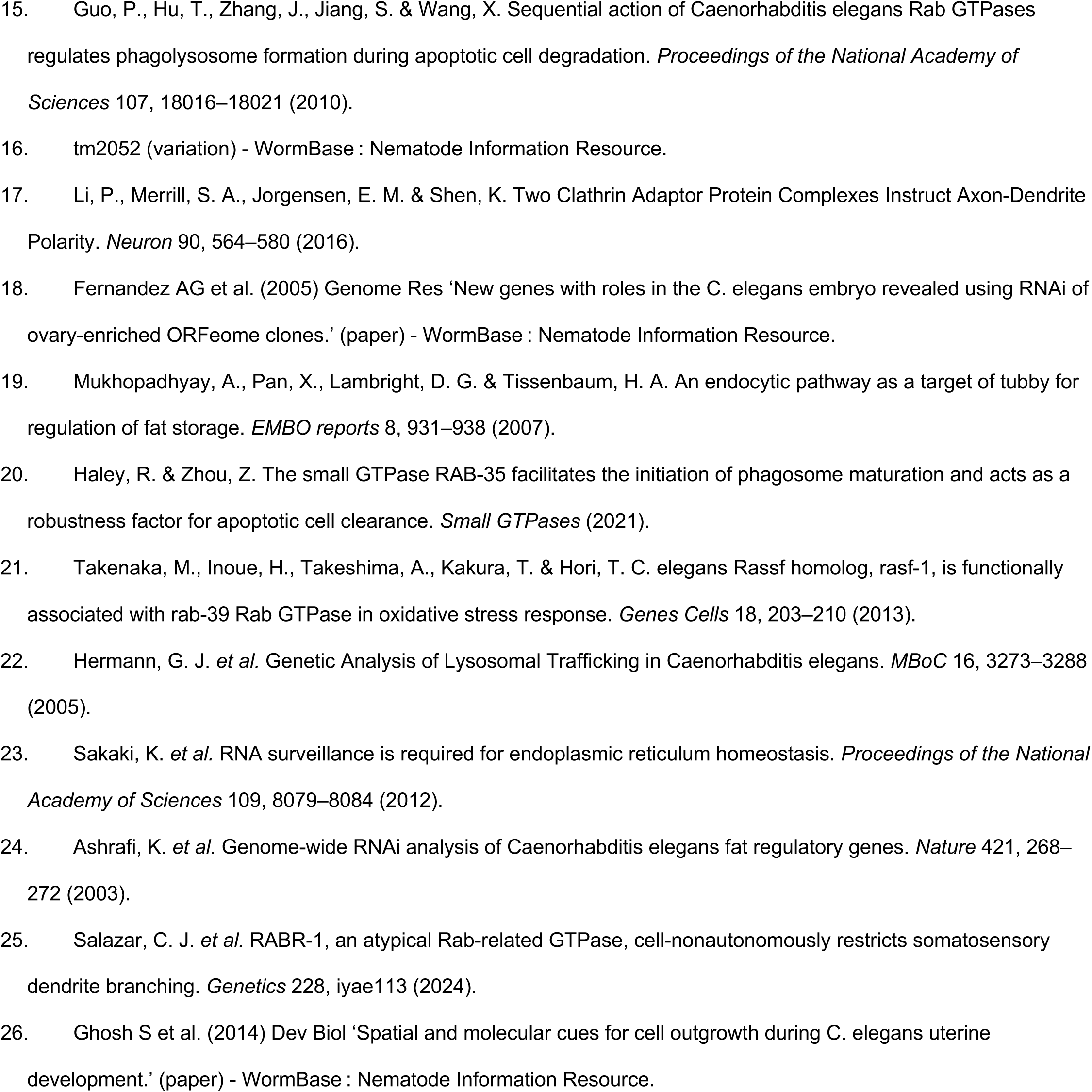
RNAi screen of all identified *C. elegans* rab-GTPase genes in a strain with markers for the germline and somatic gonad.

We probed the roles of Rab family genes in hermaphrodite gonad development (Fig. 1A), as the gonad is rapidly growing and contains proliferative germ cells (Roy et al., 2016) and large somatic cells (Byrd et al., 2014; Gordon, 2020; Li et al., 2022)—suggesting a significant dependence on intracellular trafficking. The gonad is also signal-active, with crucial regulatory interactions occurring between the soma and germline throughout life (Gopal et al., 2021; Killian and Hubbard, 2005; Kimble and Crittenden, 2005) and between the migrating somatic cells and the surrounding extracellular milieu during gonadogenesis (Agarwal et al., 2022; Meighan and Schwarzbauer, 2007; Singh et al., 2024). We designed a conservative RNAi screening approach that circumvents early lethality to reveal postembryonic developmental requirements for a third of Rab genes—*rab5, rab-7*, *rab-10, rab-11.1, rab-11.2, glo-1, rab-6.2, rab-1, rab-19, rabr-2,* and *C56E6.2—*during gonad development in *C. elegans* hermaphrodites. Secondary screening of *rab-1* in tissue-specific RNAi strains reveals that *rab-1* is required cell-autonomously for proper gonad and vulval morphogenesis and non-cell-autonomously for gamete formation. Surprisingly, the stem cell niche function of the DTC is not sensitive to cell-autonomous *rab-1* knockdown.

**Figure 1.**
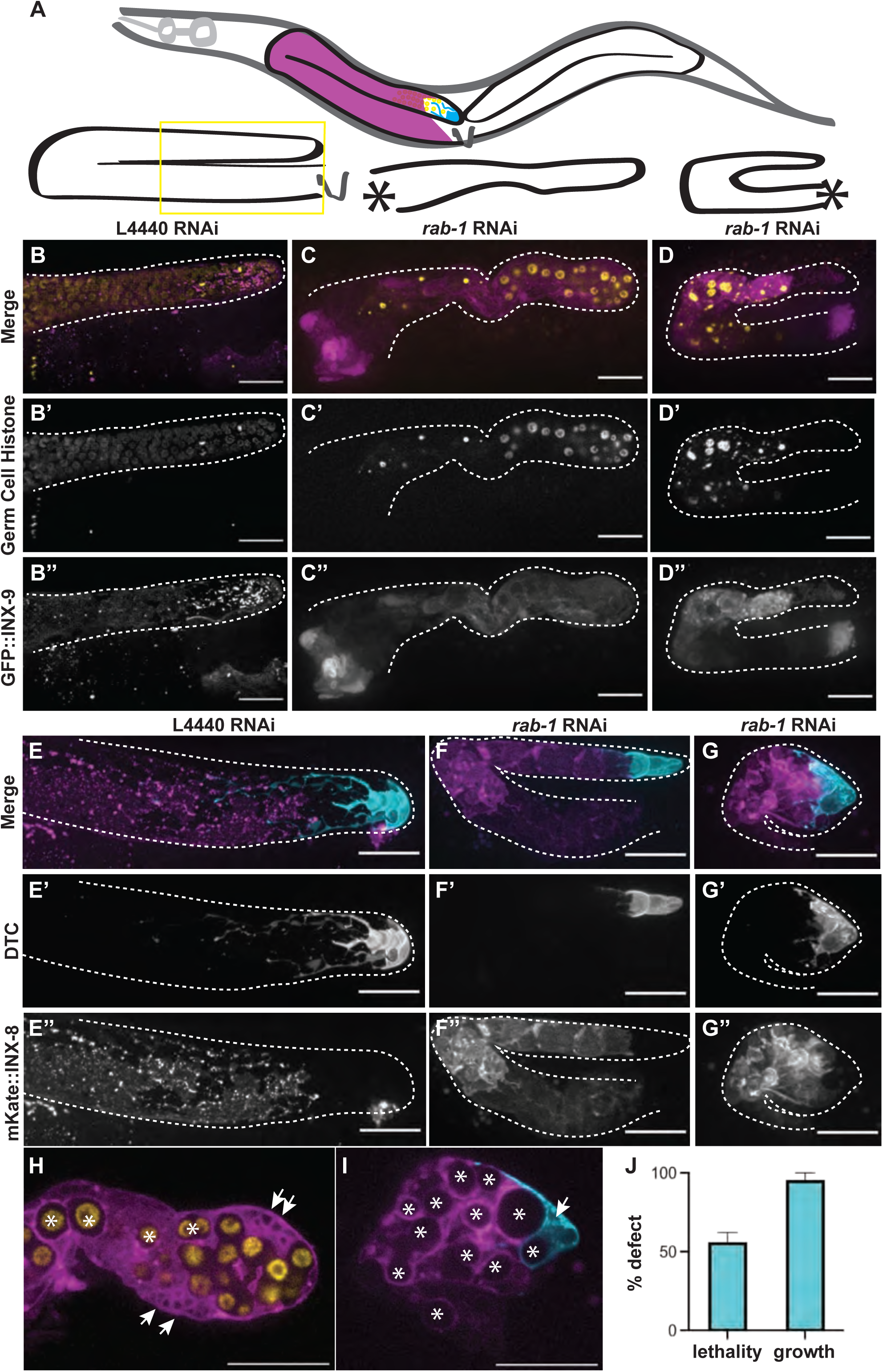
Whole-body *rab-1* knockdown leads to large-scale defects in the gonad. (A) Cartoon showing position of two gonad arms inside of worm body (above) and representative gonad shapes of control (lower left) and *rab-1* RNAi gonads (center and right). Vulva shown with V, and location where proximal gonad fails to form in RNAi samples shown with asterisks. (B-D) Representative images of germ cells (yellow; *mex-5p::H2B::mCherry::nos-2 3′UTR*) (B’-D’) and somatic gonad DTC and sheath cells (magenta; GFP::INX-9) (B’’-D’’). Adults that escaped severe embryonic and larval defects after hatching on *rab-1* RNAi (C-C’’, D-D’’) have fewer germ cells and smaller gonads in 4/16 samples compared to adults hatched on L4440 empty RNAi vector (B-B’’), and display turning and gonad growth defects (C-C”) as well as catastrophic gonad defects (D-D”). (E-G) Representative images of strain expressing markers for the DTC (cyan*; lag-2p::mNeonGreen::PLC^δPH^*) (E’-G’) and gonadal sheath cells (magenta; mKate::INX-8) (E’’-G’’). Adults that escaped severe embryonic and larval defects after hatching on *rab-1* RNAi (F-F’’,G-G’’) have DTC and sheath cells in the correct relative positions, but with aberrant morphology in 6/12 samples compared to L4440 treated controls (E-E’’). Phenotypes range from growth defects (F-F”) to catastrophic gonad defects (G-G”). (H-I) Single z-slices of images shown in C and G. Arrows show intracellular membrane protein bubbles and asterisks show encapsulation of germ cells by somatic gonad. In B and E, only the distal portion of the gonad is captured in the field of view (see yellow box in A). Gonads outlined in dashed lines. Scale bars = 20 μm. (J) Graph showing percentage lethality and percentage growth defects in larvae exposed to *rab-1* RNAi from the L1 stage. Error bars represent standard error of the mean (SEM).

## RESULTS AND DISCUSSION

### Several Rab genes are required for the development of normal gonad morphology

We took an unbiased approach to characterizing roles for Rab GTPases in *C. elegans* gonad development with a whole-body postembryonic RNAi screen in strains co-expressing a marker of germline nuclei and GFP::INX-9, a marker of the DTC and somatic gonadal sheath (Gordon et al., 2020; Li et al., 2022). Because gonad development is a postembryonic process that culminates in reproductive adulthood, and there is documented lethality caused by loss-of-function for nearly half of *C. elegans* Rab genes (Table 1), we opted not to use a strain that is sensitized for whole-body RNAi, and in some cases opted for larval exposure rather than maternal. This makes our screen conservative, and hopefully enriches for phenotypes caused by the loss of later-acting phases of Rab gene activity. We tested all 31 genes in the Rab family reported in a comprehensive phylogenetic analysis (Gallegos et al., 2012). We validated clones from the Ahringer RNAi library (Kamath et al., 2003) for 25 Rab-encoding genes, the Vidal Unique library (Rual et al., 2004) for three genes (*rab-35*, *rab-19,* and C56E6.2), and generated our own clones for three genes (*glo-1, rab-14,* and Y71H2AM.12, Table S1). Results from the screen are shown in Table 1, along with a summary of the most severe loss of function phenotype for each gene reported on Wormbase (Sternberg et al., 2024).

By analyzing knockdown phenotypes, we confirmed that loss of function of *rab-5* (Pushpa et al., 2021)*, rab-7* (Guo et al., 2010), and *rab-10* (Shi et al., 2010) cause gonad defects and identified gonad phenotypes following knockdown of *rab-11.1, rab-11.2, glo-1, rab-6.2,* and *rab-1,* as well as three genes for which little functional data is currently available: *rab-19, rabr-2, C56E6.2*.

The Rab family genes that we found to cause gonad growth defects act in a range of processes. Only two, *rab-5* and *rab-7,* are previously known to act in the cells of the gonad. RAB-5 interacts with the PAR and exocyst complexes in the germline to control levels of GLP-1/Notch at the membrane that acts as the receptor of the DTC-expressed stemness cue LAG-2 (Pushpa et al., 2021). Evidence from *Drosophila* and mammalian cells indicates that RAB7 and RAB8 are required for proper localization of NOTCH1-GFP (Court et al., 2017). In *C. elegans,* RAB-5 also acts with other Rab family members during engulfment of apoptotic cells; RAB-14, UNC-108/RAB-2, and RAB-7 follow RAB-5 recruitment and act sequentially in the formation of phagolysosomes in engulfing cells, including the gonadal sheath cells that engulf apoptotic germ cells (Guo et al., 2010). Knockdown of a RAB-7 guanine nucleotide exchange factor (GEF), *vps-45*, also causes defects in apoptotic germ cell engulfment (Kinchen et al., 2008).

Several of the Rab genes that cause gonad defects after RNAi have known roles in the *C. elegans* intestine: *rab-10, rab-11.1, rab-11.2*, and *glo-1*. The exocyst complex interacts with RAB-11 and RAB-10 in basolateral recycling and endosomal trafficking in the intestine (Chen et al., 2014; Li et al., 2024; Shi et al., 2010). The gene *glo-1* is required for the formation of lysosome-related organelles called gut granules (Hermann et al., 2005; Morris et al., 2018).

Since the gonad is exquisitely sensitive to nutrient state (Templeman and Murphy, 2018), non-cell-autonomous gonad defects could be caused by breakdown in the gut-gonad trafficking and signaling axes. While some of these genes are also expressed in neurons (e.g. *rab-10,* Zou et al., 2015), RNAi is notoriously inefficient in neurons (Calixto et al., 2010), so we conclude that neuronal knockdown is unlikely to be the cause of the defects that we observe.

Previous studies report that *rab-8* (Cram et al., 2006) and *rab-6.2* (Shafaq-Zadah et al., 2015) RNAi cause defects in later phases of DTC migration. Our screen did not detect these migration phenotypes but did detect a gonad growth defect after *rab-6.2* RNAi. RNAi is inherently variable in its efficiency, so we do not consider this negative result to be in conflict with previous findings that these genes are required for DTC migration. Indeed, they suggest that our screen is conservative, as designed.

We also observed gonad growth defects after knockdown of three genes about which little is known: *rab-19, rabr-2,* and C56E6.2*. rab-19* is most closely related to human RAB19 and RAB43, and *rabr-2*/4R39.2 is homologous to human RAB44 (Gallegos et al., 2012). C56E6.2 does not have a clear human ortholog, but it has been reported to be transcribed in the somatic gonad precursors, Z1 and Z4 (Kroetz and Zarkower, 2015). Finally, we found that normal gonad growth requires *rab-1*, the *C. elegans* paralog of human RAB1A and yeast YTP1, which is the founding member of the Rab family and key regulator of ER-Golgi transport.

### *rab-1* knockdown has profound effects on the germline that are not germ-cell-autonomous

The candidate we chose to pursue further is *rab-1*. We found that *rab-1* is important for gonad development, with *rab-1* RNAi causing severe defects in worms that survived to adulthood (Fig. 1A-I). Maternal *rab-1* RNAi caused a 97% embryonic lethality as compared to controls (control progeny n=275, *rab-1* RNAi progeny n=8 larvae by day 3 after placing L4 mothers on plates and allowing them to lay). Most worms treated from the L1 stage with *rab-1* RNAi exhibited larval growth arrest or lethality by the L3 stage (Fig. 1J), which is to be expected given the early homozygous lethality of the balanced *rab-1(ok3750)* deletion allele (Consortium, 2012). When we consider only worms that progressed through larval development after whole-body maternal *rab-1* RNAi, gonads were small, misshapen, and had very few germ cells (Fig. 1B-C). By screening in a genetic background expressing a marker of the somatic gonadal sheath and DTC (a tagged innexin protein, GFP::INX-9 (Gordon et al., 2020; Li et al., 2022), we can additionally observe that punctate membrane localization of INX-9 is impaired after *rab-1* RNAi (Fig. 1B”, C”). A similar disruption of membrane protein signal was observed after *rab-1* RNAi in a genetic background expressing another tagged innexin, mKate::INX-8 (Gordon et al., 2020; Li et al., 2022) (Fig. 1E-F). These tagged innexins and a DTC-expressed membrane-localized mNeonGreen::PLC^δPH^ reveal abnormal intracellular membranous vesicles in the DTC and sheath after *rab-1* RNAi (Fig.1H-I). While the DTC and somatic gonadal sheath are at a minimum present and in the correct relative positions—with the DTC at the tip and the gonadal sheath surrounding the germline—they have abnormal sizes, shapes, and membrane protein localization after *rab-1* RNAi.

Rab1 regulates ER-Golgi trafficking generally (Plutner et al., 1991). However, in *Drosophila* clonal analysis (Charng et al., 2014) and S2 cells (Wang et al., 2010) Rab1 has been found to play more nuanced regulatory roles, including regulating Notch and integrin signaling. Such functions have been challenging to study in genetic loss-of-function mutants due to the critical role of Rab1 in basic cell function. Performing *in vivo* studies of this crucial gene in a developmental context can expand our understanding of how a highly conserved regulator of cellular processes can nonetheless play specific developmental roles.

To elucidate the tissue-specific functions of *rab-1*, we knocked down *rab-1* predominantly in the germline in a strain bearing a *rrf-1(pk1417)* mutation for a somatic RNA-directed RNA polymerase; this strain has RNAi efficacy in the germline and some residual RNAi activity in the soma, notably the gut (Kumsta and Hansen, 2012). These animals develop mostly morphologically normal gonads (∼10% migration defect) and are smaller than age-matched controls (Fig. 2A-D), potentially due to *rab-1* knockdown outside the germline. However, embryogenesis was notably impaired after germline-specific *rab-1* RNAi, with only 1/10 of animals having superficially normal embryos in the uterus and 7/10 lacking embryos entirely (Fig. 2E). This phenotype may derive from aberrant trafficking of caveolin/CAV-1 after *rab-1* knockdown, a known role of *rab-1* in the germline (Sato et al., 2006). We determined that *rab-1* is required in the germline to produce healthy embryos, but most importantly we can conclude that loss of germline-specific *rab-1* function does not drive the dramatic gonad growth defects we see with whole-body *rab-1* knockdown (compare Fig. 2D with Fig. 1).

**Figure 2.**
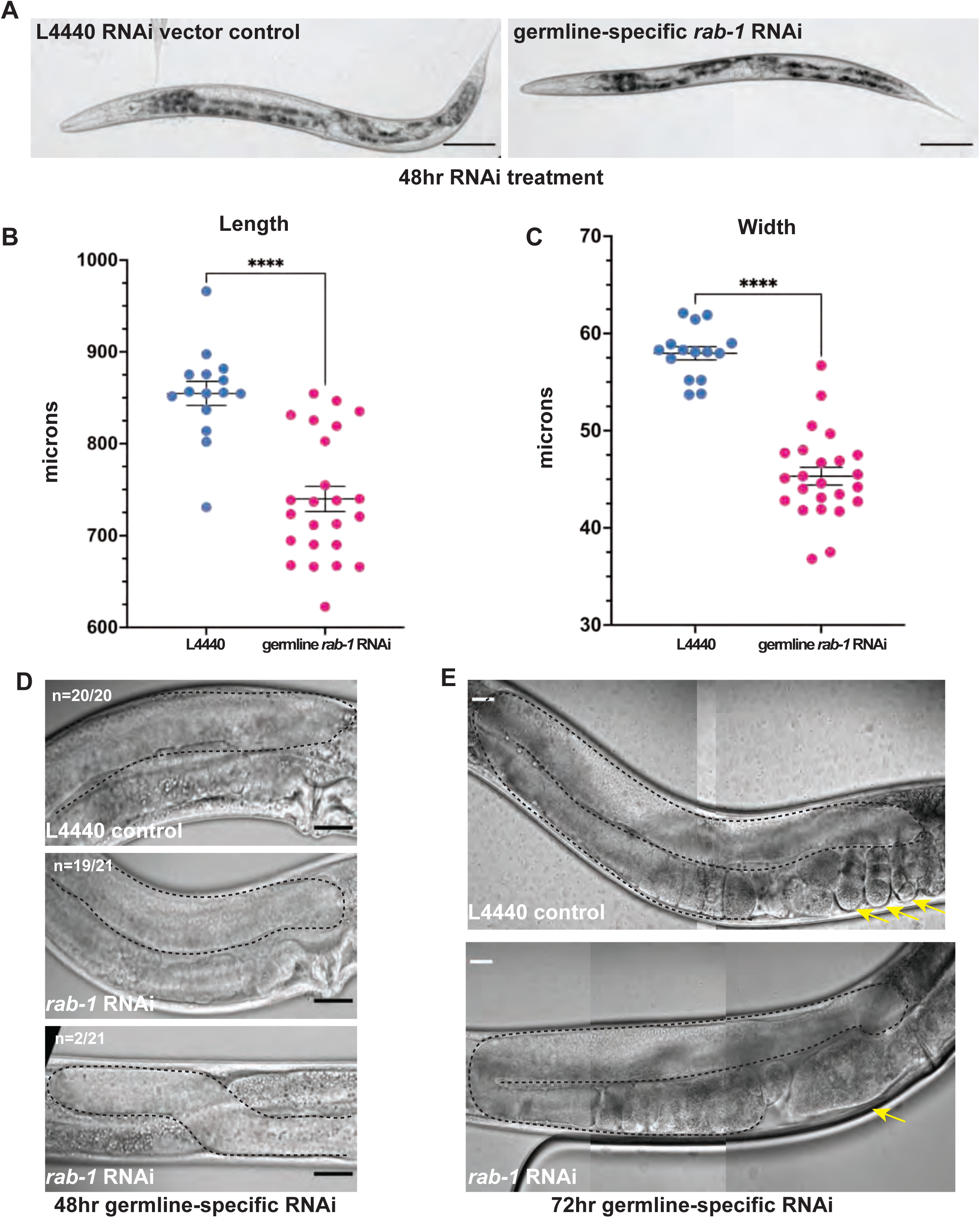
*C. elegans* with germline-specific *rab-1* RNAi knockdown are smaller, but develop largely normal gonads that produce defective embryos. (A, D-E) Representative DIC images of young adult MAH23 worms treated with L4440 control or *rab-1* RNAi for 48 (A,D) or 72 hours (E). (A, scale bar = 100 μm; D-E, scale bar = 20 μm). (B-C) MAH23 worms treated with *rab-1* RNAi (n=24) RNAi have decreased body length (B) and width (C) compared to L4440 treated controls (n=15). (D) A majority (n = 19/21) of *rab-1* RNAi treated worms (middle) developed gonads normally with 2/21 having second turn migration defects (bottom). (E) *rab-1* RNAi treated (bottom) worms have defective embryogenesis compared to L4440 (top) treated controls. Statistical significance was calculated by unpaired, two-tail Student’s t-tests, error bars represent ±SEM, and **** denotes *p*-value < 0.0001. Yellow arrows in E denote uterine contents: embryos in the case of L4440 control and abnormal masses in the case of *rab-1* RNAi. Gonads outlined in dashed lines (D-E).

### *rab-1* RNAi knockdown in *lag-2p-*expressing somatic cells of the developing reproductive system affects gonad migration and growth, as well as uterus and vulva development

Since germline knockdown of *rab-1* does not recapitulate the gonad defects of whole-body *rab-1* RNAi knockdown, we hypothesized that *rab-1* may be required in the DTC for it to function as a germline stem cell niche and as the leader cell of gonad organogenesis. We performed *rab-1* knockdown in a strain considered to have DTC-specific RNAi activity. The strain carries an *rrf- 3(pk1426)* RNAi-sensitizing mutation and an *rde-1(ne219)* loss-of-function mutation rescued by a *lag-2p::mNG::PLC^δPH^::F2A::rde-1* transgene restoring *rde-1-*dependent RNAi activity in sites of *lag-2* promoter expression, most notably the DTC, along with membrane fluorescence (Linden et al., 2017). Forty-eight hours after L1 exposure at 20°C on *rab-1* RNAi, DTC migration defects were seen in more than half of L4 larval animals (Fig. 3A-B, D). These defects all involved the arrest of gonad elongation, often accompanied by failure to turn or misdirected turning, all defects that were absent in controls (Fig. 3A-B, D). DTC migration requires signaling and adhesion, and pro-proliferative signaling from the DTC to the germline, which provides the pushing force of migration (Agarwal et al., 2022). These DTC functions are mediated by cell membrane-bound receptors, and our results suggest they may be regulated by *rab-1*.

**Figure 3.**
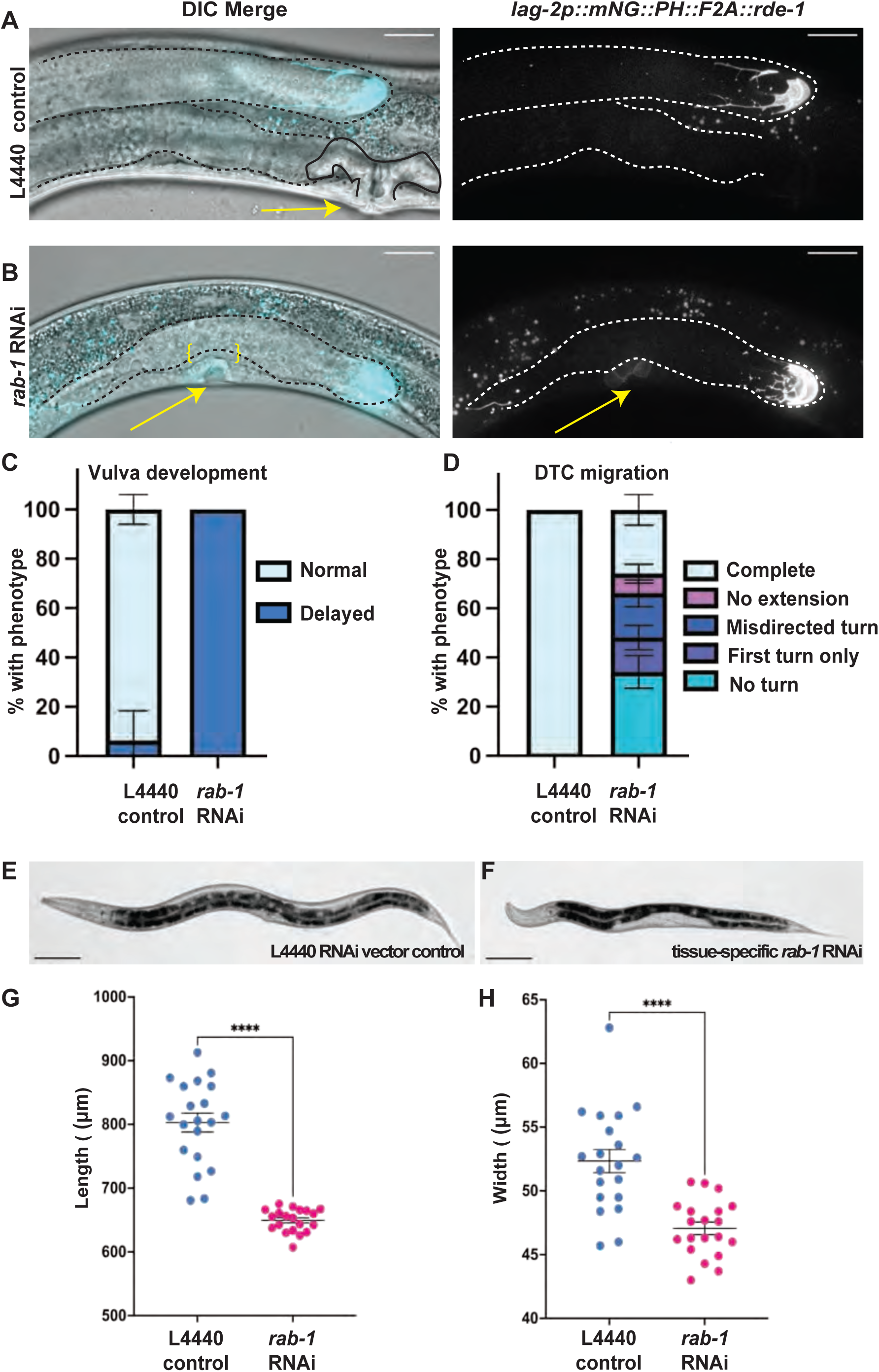
Somatic gonad-specific RNAi knockdown of *rab-1* causes gonad growth and migration, vulva, and body size defects by early adulthood. Representative images of young adult *rde-1(ne219)* mutants rescued with a *lag2p::mNG::PLC^δPH^::F2A::rde-1* transgene restoring RNAi function and driving membrane-localized fluorescence protein mNeonGreen in cells that express the *lag-2* promoter on empty L4440 vector control (A) and *rab-1* RNAi (B) for 48 hours after L1 arrest. Fluorescence merged with a single DIC z-slice (left), fluorescence alone (right, 8 μm maximum intensity projection displayed on a log scale to show DTC and dimmer vulval precursor cells, arrows). Scale bars = 20 μm. Gonads outlined in dashed lines, control uterine lumen outlined in solid black line. Unbroken basement membrane in *rab-1* RNAi treated animal bracketed in yellow. (C) Tissue-specific *rab-1* RNAi treated worms (n=17) have delayed vulva formation compared to controls (n=16). For controls, 1/16 had not yet completed vulva formation, but 17/17 *rab-1* RNAi treated worms had a delay. (D) Tissue-specific *rab-1* RNAi treated worms (n=50) have defective DTC migration compared to controls (n=31). One gonad arm scored per worm. Growth of both gonad arms was typically affected, but the deeper DTC under the gut was difficult to score for orientation of turning. All control DTCs completed migration. For *rab-1* RNAi, 17/50 had no turn, 7/50 had just a first turn, 9/50 had a misdirected second turn, 4/50 had no extension after the second turn, and 13/50 completed migration. Error bars show Standard Error of the sample proportion in C and D (see Methods). (E-F) Representative 20x brightfield images of young adult control (left) and *rab-1* RNAi (right) worms; scale bars = 100 μm). (G-H) Worms treated with *rab-1* RNAi (n=20) have decreased body length (G) and width (H) compared to controls (n=20). Mean ± S.E.M graphed in G and H. Statistical significance was calculated by unpaired, two-tail Welch’s t-tests. Length t(21.82)=10.03, p<0.0001, width t(29.39)=5.120, p<0.0001.

Surprisingly, these worms also lacked a well-differentiated uterine lumen or vulva (Fig. 3A-C). In wild-type worms, the vulva is patterned and connects to the uterus through a well-studied series of inductive events and the invasion of the anchor cell (AC) through the uterine and vulval basement membranes during the L3 larval stage (Ihara et al., 2011; Katz et al., 1995; Matus et al., 2010; Morrissey et al., 2014; Sherwood and Sternberg, 2003). In L4 worms after 48 hours on *rab-1* RNAi, we observe failure of AC invasion, with an intact basement membrane visible with Differential Interference Contrast microscopy (DIC) separating the uterus from the cells that should have formed the vulva (n = 15/17) and comparatively weak mNeonGreen expression in these vulval precursor cells (VPCs, Fig. 3B). VPCs are known to express *lag-2* (Zhang and Greenwald, 2011); expression of the *lag-2p::mNG::PLC^δPH^::F2A::rde-1* transgene that restores RNAi function is an average of ∼50x weaker in these VPCs than in the DTC at this stage, based on quantification of mNeonGreen expression (n= 16 L4 worms on *rab-1* RNAi with both vulval region and DTC captured). We see a delay in vulva formation in 17/17 of the *rab-1* RNAi-treated worms at this stage. We hypothesize that loss of *rab-1* function in the 1° vulval precursor cells prevents the completion of vulval development either through cell-autonomous defects in the 1° VPCs or the failure of these cells to signal to other cells, since VPCs engage in pro-invasive signaling to the anchor cell, (Sherwood and Sternberg, 2003) and production of DSL signaling ligands to properly induce 2° VPC fate (Chen and Greenwald, 2004).

There is no prior report of the effects of loss of *rab-1* on vulva development, but *rab-1* is required for proper development of the uterus, though its site of action in uterine development is not precisely known (Ghosh and Sternberg, 2014). The induction of both the 1° vulval precursor cell fate and the uterine pi cell fate is regulated by signaling from the anchor cell (Newman et al., 1996). The anchor cell itself arises from one of two equipotent cells (Z1.ppp and Z4.aaa) that initially express *lag-2*/DSL ligand and *lin-12*/Notch. Via lateral inhibition between the two, *lag-2* expression increases in one cell, which becomes the anchor cell; the *lin-12-*expressing cell becomes the ventral uterine (VU) cell (Seydoux and Greenwald, 1989). We confirmed that our *lag-2p::mNG::PLC^δPH^::F2A::rde-1* rescue transgene is transiently expressed in the anchor cell, but not in the other uterine cells (Fig. S1), meaning that in addition to early, continuous, and strong expression rescuing RNAi activity in the DTCs and weak expression in the VPCs, this strain also likely has transient RNAi activity in the anchor cell. Later in wild-type development, the anchor cell fuses with a subset of descendent cells of the uterine pi lineage to become the uterine-seam cell (utse) (Newman et al., 1996), so it is possible that the anchor cell could carry RNAi activity via RDE-1 protein into the utse upon fusion. Alternatively, by disrupting induction of the uterine pi cell fate by the anchor cell, RNAi activity in the anchor cell could prevent proper differentiation of uterine cell types.

Just as *rab-1* RNAi in the vulva could affect pro-invasive signaling to the anchor cell, knockdown of *rab-1* in the anchor cell could also cause defects in vulval development. For example, vulval defects are observed if the anchor cell fails to induce the primary vulval fate, fails to pattern the descendants of the 1° vulval precursor cells (Wang and Sternberg, 2000), or otherwise fails to invade and connect the uterus and vulva (Sherwood and Sternberg, 2003). In a prior RNAi screen for genes acting in the anchor cell to regulate cell invasion that used a uterus-specific RNAi strain, *rab-1* was not tested (Matus et al., 2010). When we expose the uterine-specific RNAi strain to *rab-1* RNAi from the L1 stage, we find high penetrance of uterine defects (n=9/10), no vulvaless worms, and few (n=2/10) with protruding vulvas. On the other hand, whole-body RNAi exposure initiated after anchor cell invasion begins but before major events of vulval morphogenesis (timing based on (Costa et al., 2023; Schindler and Sherwood, 2013)) showed high penetrance of vulval morphogenesis defects (n=8/8 vulval morphogenesis defect, delay, or explode through vulva). We conclude that severe defects in vulval morphogenesis are probably caused by *rab-1* RNAi in the vulval cells themselves, which express the *lag- 2p::mNG::PLC^δPH^::F2A::rde-1* RNAi rescue gene (Fig. S1).

### Prolonged tissue-specific *rab-1* RNAi knockdown in somatic gonad cells impedes vulva formation, DTC niche maturation, and germ cell proliferation

Worms with RNAi activity in *lag-2* promoter-expressing somatic cells treated with *rab-1* RNAi were also smaller than those receiving control RNAi empty vector treatment (Fig. 3D-G). We next asked whether development was simply delayed (as whole-body *rab-1* RNAi is documented to cause developmental delay, (Ghosh & Sternberg, 2014), or whether development would fail to progress with prolonged *rab-1* RNAi treatment. We allowed animals to continue to develop on *rab-1* RNAi until the age-matched controls had reached reproductive adulthood (72 hours after being released from L1 arrest at 20° C).

The penetrance of vulval morphology defects remained high after prolonged cell-specific *rab-1* knockdown, though they progressed from the “delay” phenotypes in which vulval precursor cells expressing *lag-2p::mNG* could be easily identified in adults to phenotypes like protruding vulva (pvl) or missing vulva/vulvaless (vul) (Fig. 4A-C, E).

**Figure 4.**
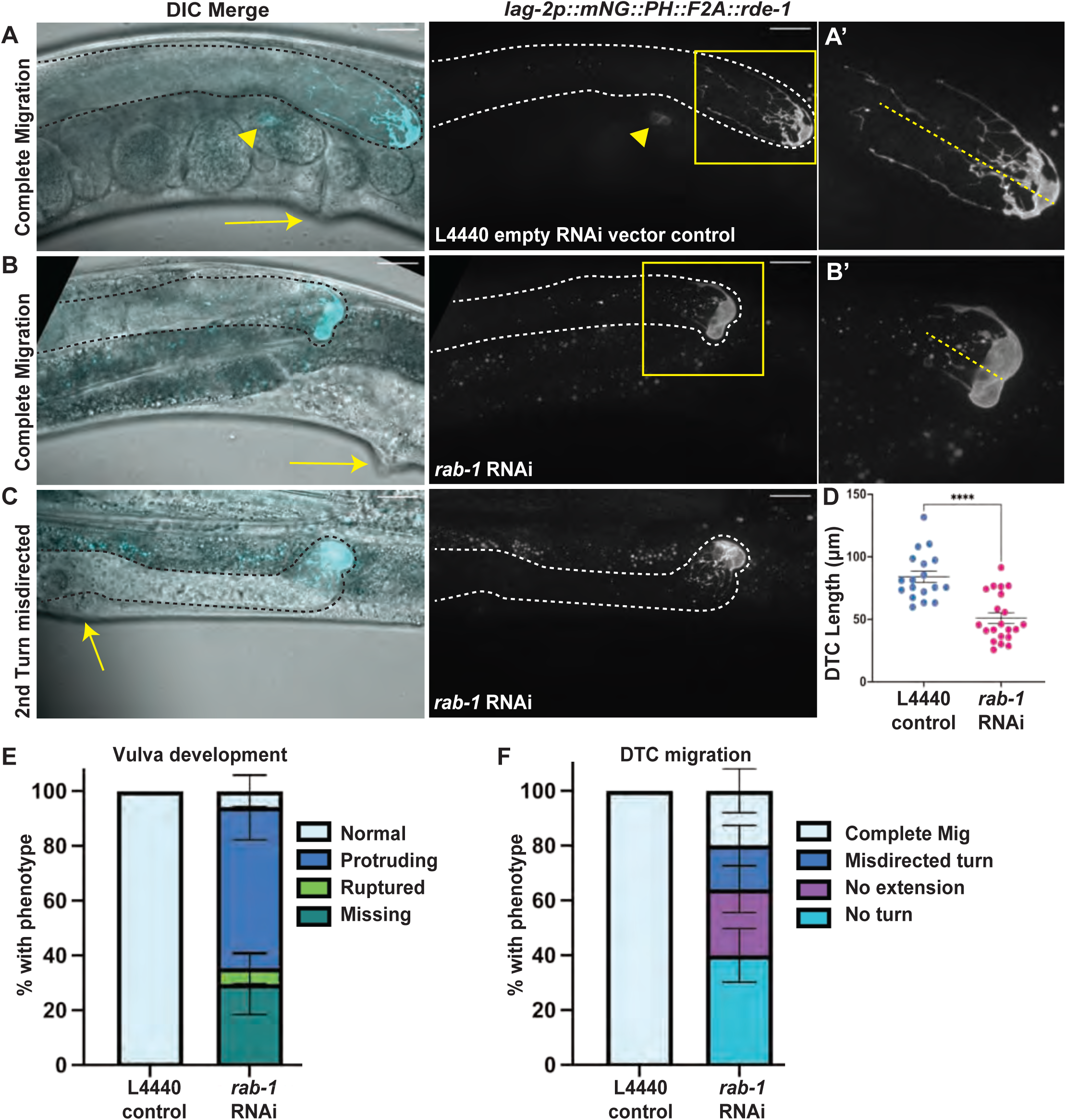
Reproductive system defects persist after prolonged *rab-1* RNAi knockdown in specific cells. Representative images (with gonads outlined in dashed black or white lines) of reproductive-age animals with tissue-specific RNAi in cells expressing a *lag2p::mNG::PLC^δPH^::F2A::rde-1* transgene restoring RNAi function and driving membrane-localized fluorescence protein mNeonGreen on empty L4440 vector control (A) and *rab-1* RNAi (B-C) for 72 hours after L1 arrest. Fluorescence merged with a single DIC z-slice (left), fluorescence alone (middle, maximum intensity projection through slices with mNeonGreen signal). A’ and B’ show insets indicated by yellow boxes. Scale bars = 20 μm. Gonads outlined in black or white dashed lines. Dashed yellow lines in A’ and B’ indicates length of DTC as measured for (D). Arrows (A-C) indicate site of expected vulval formation, showing normal vulva (A), protruding vulva (B) and vulvaless (C) phenotypes. Arrowhead in A indicates embryonic mNG expression. (D-F) Quantification of *rab-1* tissue-specific RNAi treated worm defects in DTC growth (D), vulva formation (E), and DTC migration (F) compared to controls. Both gonad arms in the same worm were scored if both were visible. (D) *rab-1* RNAi treated worms (n=21) have shorter DTCs than controls (n=18). Welch’s t-test t(36.11)=5.368, p<0.0001, error bars show S.E.M. (E) While 6/6 control samples had normal vulvas, only 1/17 *rab-1* RNAi samples had normal vulva, 5/17 had a missing vulva, 10/17 had a protruding vulva, and 1/17 ruptured through its protruding vulva on the slide. Error bars show standard error of the sample proportion (see Methods). (F) While 18/18 control samples had complete DTC migration, only 5/25 *rab-1* RNAi samples completed migration, 10/25 failed to make any turns or elongate, 4/25 made the second turn in the wrong direction, and 6/25 made both turns and then failed to extend. Error bars in E and F show standard error of the sample proportion (see Methods).

The penetrance of the DTC migration defects also remained high after 72 hours (Fig. 4A-C, F), demonstrating that gonad migration and elongation do not recover over developmental time after knockdown of *rab-1* in *lag-2* promoter-expressing cells. These results are strong evidence that *rab-1* activity in the DTC is required for gonad migration. Proper migration requires germ cell proliferation-driven gonad growth (Agarwal et al., 2022) and turning, which is regulated by several signaling pathways (Singh et al., 2024; Levy-Strumpf et al., 2015) and interactions with the basement membrane (Agarwal et al., 2022).

Upon reaching reproductive age, animals with RNAi activity in *lag-2-*expressing somatic cells also developed germline defects. Worms had smaller DTCs (Fig. 4D) and smaller germline proliferative zones (Fig. 5A-F). We also observed fewer actively dividing cells after *rab-1* RNAi, with an average of 1.67 divisions in *rab-1*-RNAi-treated worms and 3.58 dividing cells in the controls (Welch’s Two Sample t-test, t=2.6752, df=23.992, p=0.01324, 95%CI = [0.438-3.395]). However, more than half of the RNAi-treated worms had mitotic figures (Fig. 5B’), and in no case did we observe evidence of germ cell differentiation at the distal end of the gonad, which is the expected phenotype if DTC-expressed LAG-2/DSL protein is not able to signal through GLP-1/Notch receptors on the distal germ cells (Cinquin et al., 2010; Fox and Schedl, 2015). Continued maintenance of germ cells with the mitotic fate after prolonged *rab-1* RNAi suggests that localization of the stemness cue LAG-2 to the DTC membrane is not as sensitive to *rab-1* knockdown as factors regulating DTC migration behavior.

**Figure 5.**
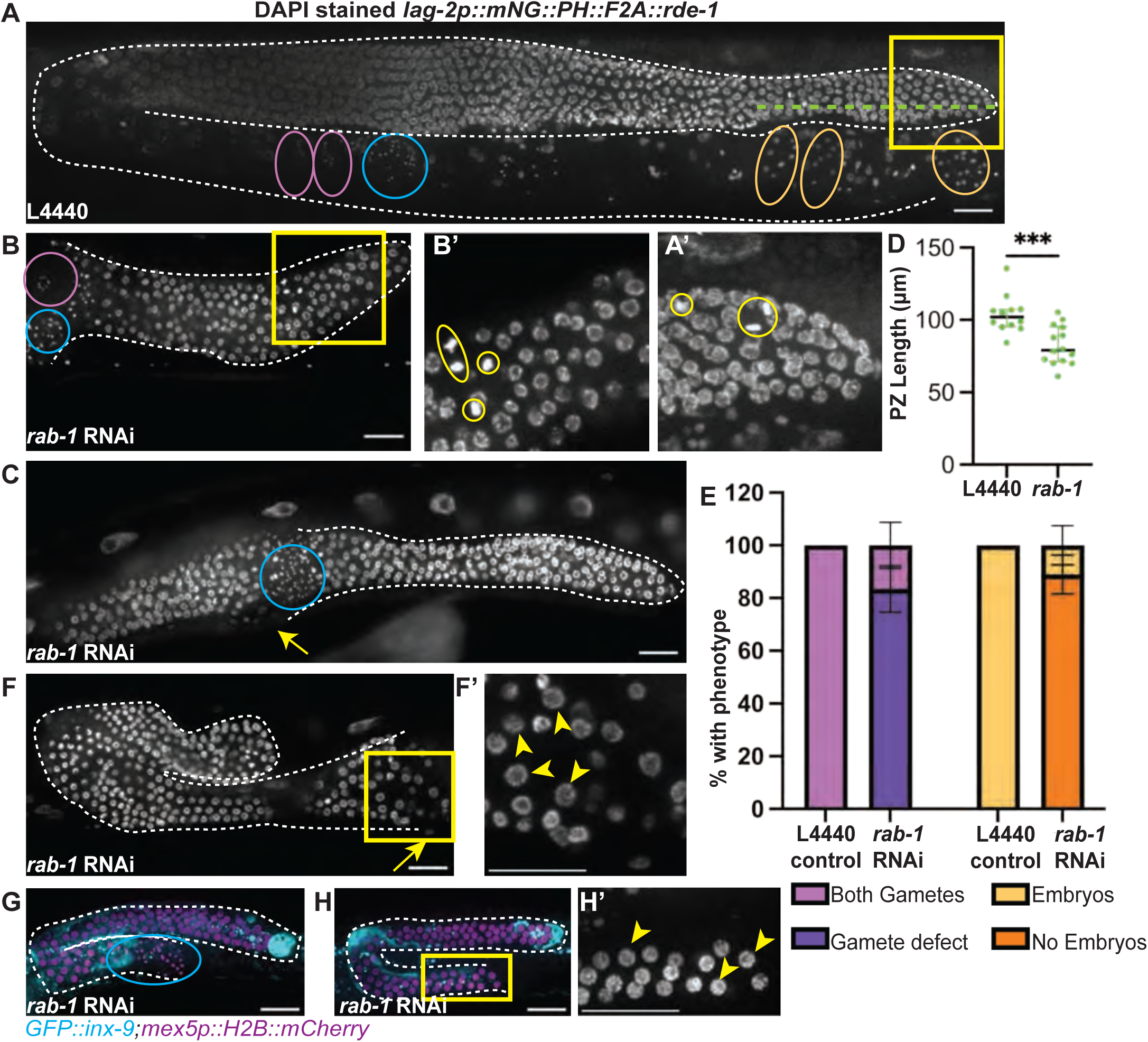
RNAi knockdown of *rab-1* in specific somatic gonad cells causes germline proliferation and differentiation defects. Representative images (with gonads outlined in dashed white lines) of DAPI-stained reproductive-age adult *rde-1(ne219)* mutants rescued with a *lag2p::mNG::PLC^δPH^::F2A::rde-1* transgene restoring RNAi function in cells that express the *lag-2* promoter on empty L4440 vector control (A-A’) and the range of defects observed after tissue-specific *rab-1* RNAi (B-C, F-F’) for 72 hours after L1 arrest. A’ and B’ show insets indicated by yellow boxes (A-B). Dividing germ cells, yellow ovals. Dashed green line shows length of progenitor zone as measured for (D). Gametes, when present, are circled in purple (oocytes) and blue (sperm). Embryos circled in orange. Arrows indicate position of expected vulva formation (C and F). Scale bars = 20 μm. (D) Tissue-specific *rab-1* RNAi treated worms (n=14) shorter progenitor zones than control (n=12), Welch’s t-test, t(23.52)=4.106, p=0.0004, error bars show median with interquartile range. (E) Gamete (left) and embryo (right) defects observed for tissue-specific *rab-1* RNAi (n=18) and control (n=12). Error bars show standard error of the sample proportion (see Methods). All 12 controls had both gametes and embryos. For *rab-1* tissue-specific RNAi, 15/18 had gamete defects (12/18 lacked gametes, 2/18 had only endomitotic oocytes, 1/18 had only sperm), 3/18 had both gametes, and 2/18 had embryos. (F-F’) A DAPI stained reproductive age animal treated with tissue specific *rab-1* RNAi with failure of gametogenesis, showing most proximal germ cells in pachytene. (G-H’) Whole-body *rab-1* RNAi treated worms expressing a somatic gonad marker in cyan (GFP::INX-9) and a marker of germ cell histones (*mex-5p::H2B::mCherry::nos-2 3′UTR)* shifted onto *rab-1* RNAi plates at the L2 stage and imaged ∼48 hours later (3 days post L1 arrest). (G) *rab-1* RNAi treated sample with sperm showing meiotic progression past pachytene. (n=4/9 RNAi treated worms) (H-H’) *rab-1* RNAi treated sample showing failure to progress through meiosis, with proximal most germ cells showing pachytene morphology (n=5/9). Yellow arrowheads indicate germ cell nuclei with pachytene morphology.

### A somatic signal that promotes pachytene exit and gamete differentiation depends on *rab-1* activity in a *lag-2*-expressing cell

Scoring the length of the proliferative zone in DAPI-stained samples revealed 4/18 samples lacking a discernible transition zone, in which germ cells in early meiotic prophase have a distinctive crescent-shaped nuclear morphology (Hubbard, 2007). The majority of animals also fail to make gametes—both sperm and eggs—normally (Fig. 5A-E, G). When a germline is shorter than normal, meiotic entry delay can result because much of the germline remains within niche signaling range of the DTC (Kimble and White, 1981). There is also a “latent niche” DSL ligand signal from the proximal gonad that can support germline mitotic fate if undifferentiated germ cells come in contact with the proximal gonad (McGovern et al., 2009). However, we did not observe proximal mitotic figures in these very small gonads (as in *pro* mutants, McGovern et al., 2009), and most samples did show evidence of meiotic entry with a well-demarcated transition zone, suggesting that germ cells are escaping the niche signal (Fig. 5B, n=14/18).

Instead, we propose that the phenotypes we observed after prolonged *rab-1* RNAi knockdown in *lag-2* promoter-expressing cells represent failure to progress through meiotic pachytene (Church et al., 1995; Lee et al., 2007; McCarter et al., 1997). Of the samples that lacked differentiated gametes entirely, half showed DAPI signal consistent with meiotic pachytene in the proximal-most germ cells (n=5/10, Fig. 5F-F’). Late L2 exposure to whole body RNAi using a strain marking germ cell histones (Fig. 5G-H’) also resulted in the most proximal germ cells having pachytene morphology (n=5/9, Fig. 5H-H’), with the remaining samples showing nuclear morphology representing later stages of male gamete meiosis (Shakes et al., 2009) or mature sperm (n=4/9, Fig. 5G) at the most proximal region of the gonad. Oocytes were never observed (n=0/9).

A defect of pachytene exit affecting both spermatogenesis and oogenesis with similar penetrance was observed in a prior study after laser ablation of both sheath-spermathecal progenitor cells in a gonad arm (57% of gonads had sperm, only 4% of arms made a single oocyte, (McCarter et al. 1997). The genetic regulation of meiotic cell cycle progression is complex, especially for the oogenic germline (Arur, 2017), in which MPK-1 (the *C. elegans* ortholog of ERK, the primary MAPK required for germline differentiation) signaling plays a crucial role (Das and Arur, 2022; Lee et al., 2007). MPK-1/MAPK signaling regulates the germline both cell-autonomously (Lee et al., 2007) and non-cell-autonomously (Robinson-Thiewes et al., 2021). Nutritional inputs also regulate MPK-1/MAPK-mediated progress through meiosis in the oogenic germline through the insulin-like receptor encoded by *daf-2*, with neuronal insulin-like peptides proposed as the signals that activate germline DAF-2 (Lopez et al., 2013). The source and identity of the somatic gonad signal that promotes pachytene exit have not yet been identified.

We propose that by knocking down *rab-1*—and thereby a key step in the secretory pathway—in *lag-2-*expressing cells of the reproductive system, we have identified a subset of cells (DTC, sheath-spermathecal cells, anchor cell, vulval cells) within which may be the source of a somatic gonad pachytene exit signal, or at least a cell or cells required for the proper development of the source of that signal. Some *lag-2* promoter-driven transgenes that activate in Z1 and Z4, the somatic gonad progenitor cells, show residual expression in other somatic gonad cell types later in development like the somatic gonad sheath cells (Blelloch et al., 1999; Killian and Hubbard, 2005). We saw expression of the *lag-2p::mNG::F2A::rde-1* rescue transgene in Z1 and Z4, and then in the sheath-spermathecal progenitor (SS) cells for a brief time in the L2 larval stage (Fig. S1). The SS lineage is the only point of convergence between *lag-2p*+ cells and cells targeted by the SS ablation experiment that blocked pachytene progression (McCarter et al., 1997), though the *lag-2* promoter is active in the SS cells for a very brief time. Further work on the sensitive developmental window in which *rab-1* RNAi in *lag-2p*+ cells inhibits pachytene progression will further narrow down the source of a somatic signal necessary for gamete differentiation.

### Conclusions and Future Directions

We found that eleven of the 31 Rab GTPase-encoding genes in *C. elegans* play a role in gonad development and that *rab-1* regulates development of the somatic gonad and germline in both cell-autonomous and non-cell-autonomous ways. Neither germline-specific *rab-1* RNAi nor *rab-1* RNAi in *lag-2* promoter-positive cells recapitulated the catastrophic gonad and germline defects we observed after whole-body *rab-1* RNAi treatment (Fig. 1), so we conclude that systemic *rab-1* is essential for germline and gonad growth. In the future, intestinal *rab-1* should be examined for a role in gonad development, as several known intestine-expressed Rab genes (*rab-10, rab-11.1, rab-11.2*, and *glo-1*) also caused gonad defects when knocked down with whole-body RNAi (Table 1). Tissue-specific RNAi implicates cell-autonomous *rab-1* in the formation of normal embryos, vulva morphogenesis, the proper development of uterine cells, and DTC migration. The germline requires non-cell-autonomous *rab-1* expression for normal proliferation and for pachytene exit. Surprisingly, the least-sensitive feature of the reproductive system to *rab-1* RNAi knockdown is the stem cell niche function of the DTC, which survives even strong knockdown of *rab-1* in the DTC. This study motivates future investigations into the role of *rab-1*-independent signaling from the stem cell niche to the germline, as well as *rab-1*-mediated signaling to promote pachytene exit.

## METHODS

Sections of this text are adapted from (Li et al., 2022), as they describe our standard laboratory practices and equipment.

### EXPERIMENTAL RNAi

A single colony of *E. coli* HT115(DE3) containing the L4440 plasmid with or without (as a control) a dsRNA trigger insert from the Ahringer (Kamath et al., 2003) or Vidal (Rual et al., 2004) RNAi libraries, or our own clone in the case of *rab-14, glo-1* and Y71H2AM.12, was grown as an overnight culture containing ampicillin (100 μg/ml, VWR (Avantor), Catalog no. 69-52-3) at 37°C. Expression was induced with 1mM IPTG (Apex BioResearch Products, cat# 20-109) for one hour at 37°C, and 150-300 µl of induced RNAi culture was plated on NGM plates and allowed to grow on the benchtop at least overnight. Glycerol stocks were prepared from the pre-induction overnight culture for storage at -80°C and future use, and a subsample was miniprepped and sent for sequencing to verify sequence of insert.

Worm populations were synchronized by bleaching according to a standard egg prep protocol (Stiernagle, 2006), plated on NGM plates seeded with RNAi-expressing bacteria as arrested L1 larvae, and kept on RNAi until the time of imaging.

### Initial Rab Whole-body RNAi Screening

Worms were maintained at 20°C. In the case of maternal RNAi exposure, L4 hermaphrodites with somatic gonad membrane protein and germline nuclear markers (see Strains section) were added to RNAi plates and allowed to feed on RNAi-expressing bacteria prior to egg laying. In the case of L1 RNAi exposure, synced L1s were plated on RNAi plates. Offspring were assessed for adult phenotypes four days later for maternal exposure and 3 days later for L1 exposure, both at 20°C. *rab-1* RNAi treatment resulted in high levels of embryonic lethality or larval arrest. In this case, RNAi was repeated by dropping egg-prepped L1 larvae directly onto *rab-1* RNAi plates and assessing adult phenotypes three days later. Each RNAi treatment was paired with an L4440 empty vector control treatment, none of which ever showed gonad growth defects. Therefore, an RNAi treatment that caused even a single incidence of gonad growth defect was counted as a “hit”.

### Tissue-specific and temporal RNAi of *rab-1*

The same *rab-1* RNAi clone used in the initial screen was rescreened in animals with tissue-specific RNAi activity. Germline specific RNAi used a strain MAH23 carrying a mutation in *rrf-1(pk1417)*; it has some residual RNAi function in somatic cells (Kumsta and Hansen, 2012), but does not phenocopy whole-body knockdown of *rab-1* and instead shows phenotypes in the germline. A second tissue-specific strain has RNAi activity only in *lag-2p+* cells, namely the DTC, anchor cell, and primary vulval precursors; this strain, NK2115, carries an *rde-1(ne219)* loss of function that prevents RNAi activity globally, with RNAi function restored in the DTC (in an operon along with a coding sequence for membrane-tethered mNeonGreen) by a transgene *lag-2p::mNG::PLC^δPH^::F2A::rde-1* and a *rrf-3(pk1426)* mutation that enhances RNAi. DTC-specific RNAi treatment was conducted at 16°C due to a temperature-sensitive *rrf-3(pk1426)* mutation in the DTC-specific RNAi strain (Linden et al., 2017). A third tissue-specific strain, NK1316, has uterine specific RNAi activity with *fos-1ap::rde-1* (Hagedorn et al., 2009; Matus et al., 2010), and functions by restoring RDE-1 protein activity in *rde-1(ne219)* mutant animals only to those cells of the somatic gonad that are under the control of the *fos-1a* promoter, expressed in uterine cells in the mid to late L2. NK1316 also carries the *rrf-3(pk1426)* mutation that enhances RNAi.

Temporal whole-body RNAi of *rab-1* targeted two developmental windows at 20°C. To assess whether *rab-1* knockdown affects vulval morphogenesis due to loss of function in the anchor cell or the vulval cells themselves, we shifted *GFP::inx-9;mex-5pH2B::mCherry::nos-2 3′UTR* worms to *rab-1* RNAi at 26 hours post L1 arrest, meaning that RNAi-mediated gene knockdown is expected ∼4 hours later at the earliest, at which point anchor cell invasion is underway (28 h-31.5 h post L1 arrest (Costa et al., 2023). Important developmental events of vulval morphogenesis unfold slightly later, from ∼31 h-34 h (Schindler and Sherwood, 2013).To assess whether *rab-1* knockdown causes pachytene arrest, we shifted worms of the same strain to *rab-1* RNAi at 15 h post hatching in the L2 larval stage to circumvent developmental arrest/lethality we see with *rab-1* RNAi treatment beginning at L1 (Fig. 1J). Sample sizes for whole-body RNAi of *rab-1* are low due to the propensity of *rab-1* RNAi treatment to cause worms to rupture through the vulva before or during mounting on a slide.

### Scoring gonad defects

DTC migration was scored by the following categories: CT: complete migration; NT: no turn; FT: first turn only; ST: second turn complete, but no extension; MD: misdirected second turn. Vulva formation was scored as wild-type, protruding vulva (pvl), and vulvaless (vul) at 72 h, and wild-type or delayed (large, round, *lag-2p::mNG*+ VPCs still visible) at 48 h. DTC length was measured from tip to end of the longest process (Linden et al., 2017).

DAPI-stained adult worms were scored for presence of oocytes (large cells with chromosomes in diakinesis), sperm (small pinpoints of DAPI), and embryos (multicellular structures in the proximal gonad). Progenitor zone was scored from the tip to the first row of germ cells with two crescent-shaped nuclei (Hubbard, 2007). Some *rab-1* RNAi samples lacked an identifiable transition zone and were not scored. Mitotic figures were scored as a single bright metaphase plate or a pair of anaphase DAPI bodies; these were scored in the distal gonad and found to be absent in the proximal gonad.

### Confocal imaging

All images were acquired at room temperature on a Leica DMI8 with an xLIGHT V3 confocal spinning disk head (89 North) with a 63× Plan-Apochromat (1.4 NA) objective and an ORCAFusion GenIII sCMOS camera (Hamamatsu Photonics) controlled by microManager. RFPs were excited with a 555 nm laser; GFPs and mNGs were excited with a 488 nm laser. Z-stacks through the gonad were acquired with a step-size of 1 µm unless otherwise noted.

Worms were mounted on agar pads in M9 buffer with 0.01 M sodium azide (VWR (Avantor) Catalog Number 26628-22-8).

### Image analysis

Images were processed in FIJI89 (Version: 2.14.1/1.54f). Larger images tile several acquisitions of the same sample (Preibisch et al., 2009).

### Statistical analysis

Sample sizes vary slightly for measurements gathered from the same dataset if certain cell types were not clearly represented (for example, if the vulva is visible but the DTC is under the gut, or the image quality allows the DTC to be scored for position but not for length of processes). Sample sizes stated in figure legends or text reflects the number of samples analyzed for the specific feature being measured. Welch’s two-sample t-tests were used to compare *rab-1* RNAi to controls.

Standard Error of the sample proportion for the histograms in Figures 3, 4, and 5 (after Levy-Strumpf et al., 2015) was calculated using the equation, 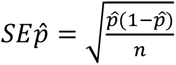where *p̂* is the percentage of specimen of the total observed *(n)* with the phenotype and error bars reflecting this calculation were added to plots using GraphPad Prism (Prism 10 for Mac OS; Version 10.1.0 (264), October 18, 2023).

## Acknowledgements

We would like to acknowledge undergraduate lab members Jayce Proctor and Kayah Takei for experimental assistance. We would like to acknowledge Dave Sherwood and Eric Hastie for helpful advice. Some strains were provided by the CGC, which is funded by NIH Office of Research Infrastructure Programs (P40 OD010440) and are to be requested directly from CGC. Funded by NIGMS grant R35GM147704 to K.L.G.

---

**Figure S1.**
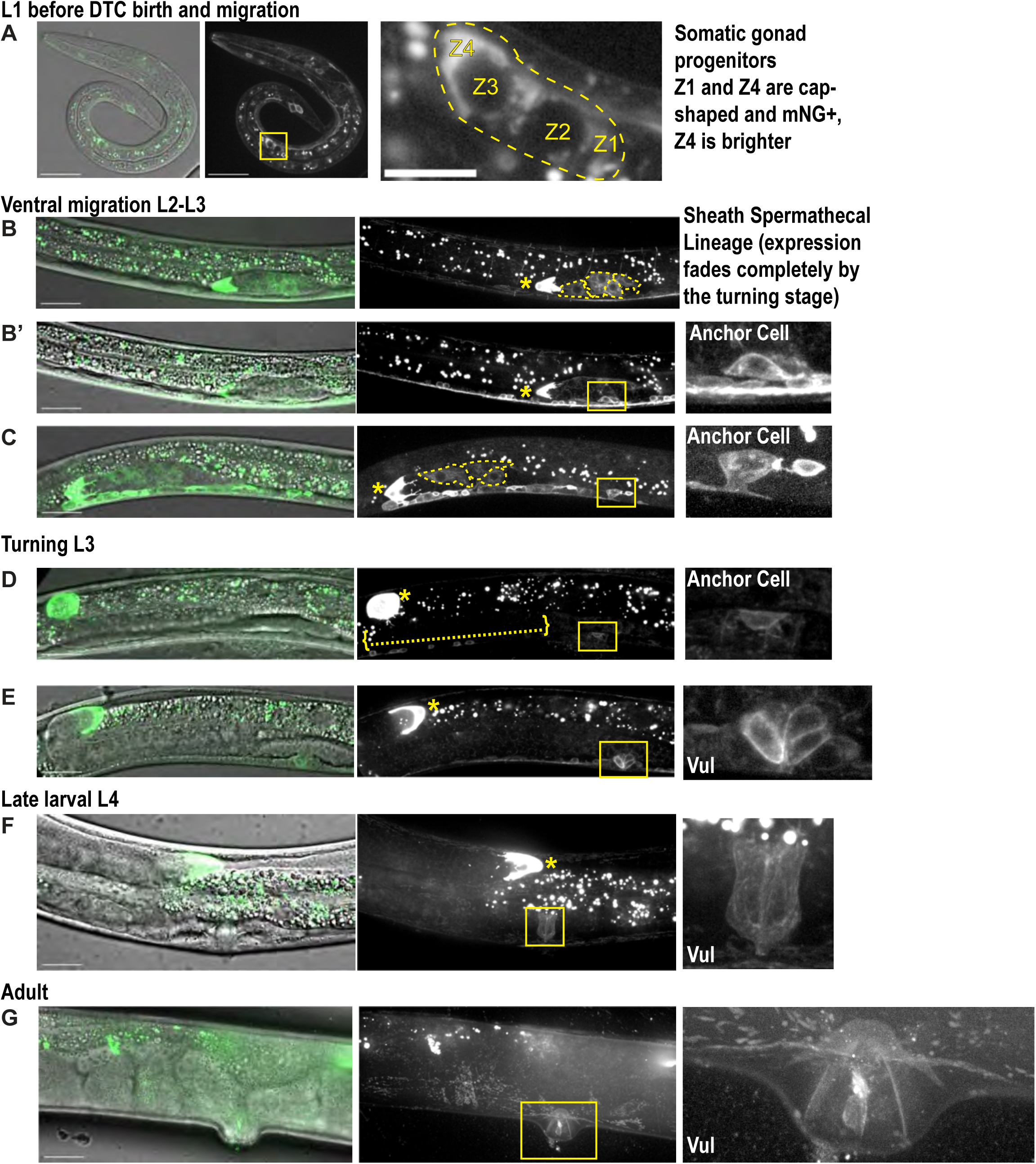
Locations of *lag2p::mNG::F2A::rde-1* rescue transgene expression during development. Representative images of the strain bearing the *lag2p::mNG::PLC^δPH^::F2A::rde-1* allele (with mutations in *rrf-3(pk1426)* and *rde-1(ne219)*) in the L1 (A), L2 (B-C), turning/late L3 (D-E), L4 (F), and Adult (G) life stages in which RNAi is presumably active in mNG+ cells (with the exception of neurons, which are not sensitive to RNAi (Calixto et al., 2010), seen along the ventral body wall in some images and in superficial circumferential projections in B). Single slices or projections through 0.5-2.5 microns from confocal z-stacks, DIC images merged with green fluorescence (right), and green fluorescence alone (grayscale, center and insets). Imaged with 0.5 micron step size, except (A), which was imaged with 0.2 micron step size. Greyscale images shown with fluorescence logarithmically scaled to enhance visibility of dim signal. Boxes show region of inset. Asterisks mark distal tip. (A) Somatic gonad progenitor cells Z1 and Z4 express mNG in L1 arrested larvae, with substantially more expression in Z4 than Z1; primordial germ cells Z2 and Z3 do not express the rescue transgene. (B) In the L2, cells of the sheath-spermathecal (SS) lineage (outlined in yellow dashed lines) are mNG+ early, possibly as residual expression from the somatic gonad precursor cells; these cells are no longer visible by the turning stage in L3 (brackets in D). (B’) Projection through deeper Z-slices of the same sample in B. The anchor cell (AC) is mNG+ from the late L2 when it is first born. (C) SS cells are still faintly mNG+ at the time of AC invasion (inset). (D) By the time of DTC turning in the L3, no SS expression is observed, and signal at the future site of vulva formation (inset) is very dim. (E) As the turn is completed, the vulval precursor cells (inset) are mNG+. (F) The vulva remains mNG+ in L4 larvae and (G) adults. Scale bars = 20 μm.

**Table S1.**
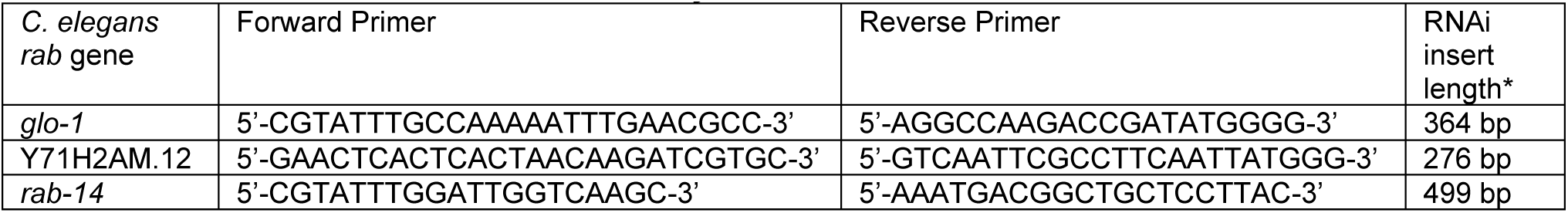
RNAi clones made for this study.

